# Body temperature regulates glucose metabolism and torpid behavior

**DOI:** 10.1101/2024.01.29.577738

**Authors:** Ming-Liang Lee, Ching-Pu Chang, Chitoku Toda, Tomomi Nemoto, Ryosuke Enoki

## Abstract

Glucose is a significant energy resource for maintaining physiological activities, including body temperature homeostasis, and glucose homeostasis is tightly regulated in mammals. Although ambient temperature tunes glucose metabolism to maintain euthermia, the significance of body temperature in metabolic regulation remains unclear owing to strict thermoregulation. Activation of Qrfp neurons in the preoptic area induced a harmless hypothermic state known as Q-neuron-induced hypothermia and hypometabolism (QIH), which is suitable for studying glucose metabolism under hypothermia. In this study, we first observed that QIH mice had hyperinsulinemia and insulin resistance. This glucose hypometabolic state was abolished by increasing the body temperature to euthermia. Moreover, QIH-mediated inappetence and locomotor inactivity were recovered in euthermia QIH mice. These results indicate that body temperature is considerably more powerful than ambient temperature in regulating glucose metabolism and behavior, and hypometabolism in QIH is secondary to hypothermia rather than modulated by Qrfp neurons.

**Highlights:**

- QIH reorganizes glucose homeostasis which is unchanged by fasting.
- QIH mice exhibit glucose hypometabolism with hyperinsuliemia and insulin resistance.
- Increased body temperature abolishes QIH-mediated hypometabolism and torpid behaviors.
- Body temperature is a strong factor in controlling metabolism and behavior.
- Body temperature-mediated glucose metabolism is reversible.

## Introduction

Glucose is the major energy source and is universally used in almost every cell in animals. Glucose metabolism is tightly regulated by insulin to maintain euglycemia. The actions of insulin are orchestrated by plasma insulin levels and tissue insulin sensitivity to regulate endogenous glucose production and tissue glucose uptake from the blood (Hatting et al., 2018; Leto and Saltiel, 2012; Norton et al., 2022). Metabolic disorders or diseases such as diabetes can disrupt glucose homeostasis, in which insulin resistance develops and hyperglycemia and hyperinsulinemia occur.

Temperature is a significant factor in the regulation of whole-body metabolism (Sadler et al., 2022; Westerterp-Plantenga et al., 2002). As experimental animals such as mice are frequently housed at 20°C–25°C, which is lower than their thermoneutral zone, a lot of energy is used for heat generation to maintain normal body temperature by increasing systemic metabolism (Neff, 2020). Conversely, mice housed in the thermoneutral zone show lower metabolic rates with lower oxygen consumption because the energy cost of thermogenesis is inhibited (Ganeshan and Chawla, 2017). Normal body temperature, euthermia, is essential for normal physiology and survival; therefore, it is strictly regulated within a narrow range. This strong body thermoregulation makes examining the relationship between body temperature and metabolism difficult. Although several chemicals such as N6-cyclohexyladenosine (CHA) or 2-deoxy-D-glucose (2DG) are used to induce temporary hypothermia, they have several off-target effects by unwanted disturbance of neuronal activity and are not suitable for investigating glucose metabolism (Alvarsson and Stanley, 2018; Ohno and Watanabe, 1996; Olson et al., 2013). Therefore, the relationship between glucose metabolism and body temperature remains unclear.

Pyroglutamylated RFamide peptide (Qrfp) is a member of the RFamide-related peptide family (Primeaux et al., 2013). This versatile hypothalamic neuropeptide regulates various physiological processes, including food intake, glucose metabolism, locomotor activity, and blood pressure (Chen et al., 2016; El-Mehdi et al., 2020; Takayasu et al., 2006; Zagorácz et al., 2015). Recently, Qrfp-expressing neurons in the preoptic area (Qrfp^POA^) were noted to induce hypothermia for days. Mice with Q-neuron–induced hypothermia and hypometabolism (QIH) have lower body temperature and decreased systemic metabolism with extremely inhibited locomotor activity and food intake without tissue damage (Takahashi et al., 2020). This suggests that QIH is a benign form of hypothermia and can be an appropriate model for studies under hypothermia. Therefore, to examine glucose metabolism under hypothermia, we used the QIH model. In this study, we observed that QIH-mediated hypothermia was accompanied by reorganized glucose homeostasis due to hyperinsulinemia and severe insulin resistance. This diabetes-like glucose hypometabolism was temperature dependent and was abolished in euthermic QIH mice. Therefore, we propose that body temperature is sufficient as a critical factor in regulating glucose metabolism and that QIH-mediated glucose hypometabolism is secondary to hypothermia rather than regulated by Qrfp^POA^.

## Results

### QIH reorganizes glucose homeostasis which resists fasting-mediated hypoglycemia

To investigate glucose metabolism under QIH, designer receptors exclusively activated by designer drugs (DREADD) was used to activate Qrfp^POA^. AAV delivering Cre-dependent hM3Dq was bilaterally injected into the POA of Qrfp-iCre mice, whereas AAV-DIO-mCherry was injected into the control group of Qrfp-iCre mice (Figure 1A). The temperature of brown adipose tissue (T_BAT_) expeditiously decreased following clozapine injection. T_BAT_ was lower than 30°C within 1 h following clozapine injection, and this QIH status was maintained for at least 24 h (Figures 1B and 1C). c-fos expression showed that Qrfp^POA^ remained highly activated 24 h after clozapine injection (Figures 1D and 1E). QIH not only induced hypothermia but also abolished appetite (Figure 1F). As the feeding state strongly affects glucose metabolism (Jensen et al., 2013), food was removed from either QIH or control mice after clozapine injection to evaluate glucose metabolism in the following experiments (Figure 1G).

**Figure 1.**
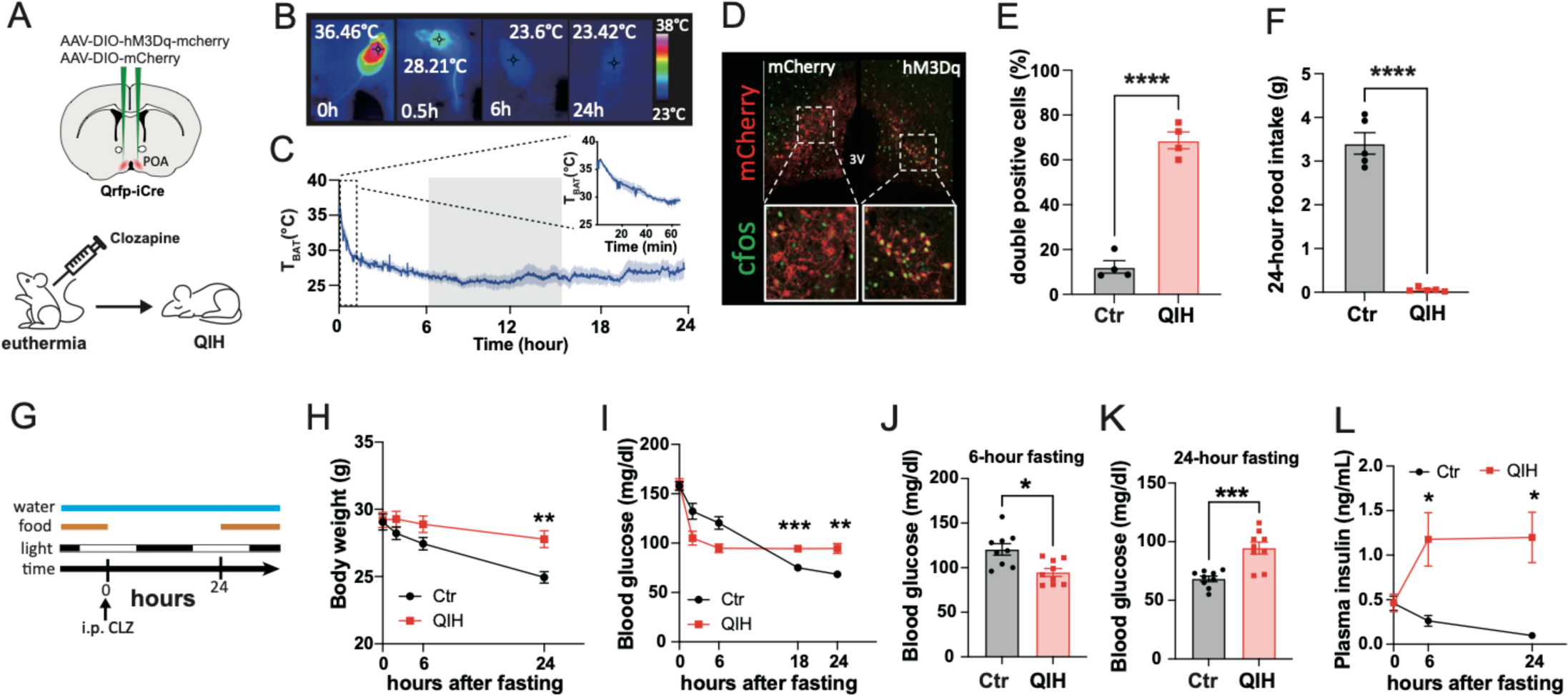
QIH reorganizes glucose homeostasis with a fasting independent property. (A) Diagram of viral injection for chemogenetics (B) Representative thermography showing BAT temperature of QIH mice at different time points (C) Tracing of T_BAT_ after clozapine injection (n = 9) (D) Representative micrographs showing immunofluorescent cFos staining 24 h after clozapine injection in the control (left) and QIH (right) groups (E) Quantification of cFos expressing Qrfp neurons in the preoptic area from the control (n = 4) and QIH (n = 4) groups (F) Twenty-four-hour food intake of control (n = 5) and QIH (n = 5) mice (G) Experimental design for evaluating glucose metabolism (H) Body weight (BW) of control (n = 9) and QIH (n = 9) mice during fasting (I–K) Blood glucose levels of control (n = 9) and QIH (n = 9) mice during fasting (I) Blood glucose levels of control and QIH mice at different time points after food deprivation (J) Comparisons of blood glucose levels in control and QIH mice after 6-h fasting (K) Comparisons of blood glucose levels in control and QIH mice after 24-h fasting (L) Plasma insulin levels in control (n = 8) and QIH (n = 8) mice during fasting All data are presented as means ± SEM; *p < 0.05; **p < 0.01; ***p < 0.001; ****p < 0.0001.

To understand the responsiveness of glucose homeostasis to changes in energy levels, we removed food at the same time as the clozapine injection and recorded Body weight (BW) and blood glucose levels. During fasting, both the control and QIH groups showed decreased BW (Figure 1H). The control mice lost up to 15% of their BW, whereas the QIH mice lost only approximately 5% of their BW after the 24-h fasting (Figures 1H and S1A). To confirm that the difference in BW loss was not caused by undigested food or feces, tissue weights were compared between the control and QIH mice. The weights of the stomach and intestine with undigested food were higher in QIH animals (approximately 0.1 g and 0.3 g, respectively) (Figures S2B and S2C). However, the total changes in the stomach and intestine were too small to explain the differences in BW loss between the control and QIH mice (2–3 g) (Figures 1H and S2A). The weights of other tissues including the skeletal muscle, BAT, epididymal white adipose tissue (eWAT), inguinal WAT (iWAT), mesenteric WAT (mesWAT), and liver were higher in the QIH group (Figures 2D–2J), suggesting that QIH conserved energy and prevented BW loss. Fasting resulted in a gradual decrease in blood glucose levels in control mice but not in QIH mice (Figures 1I and S1B). Instead, the blood glucose levels of QIH mice were stable and unaffected by fasting after a prompt decrease in blood glucose levels in the first few hours after clozapine injection (Figure 1I). The blood glucose levels of QIH mice were similar between 6- and 24-h fasting periods (Figures 1I and S1C). Conversely, the blood glucose levels of control mice continued to decrease from 6 to 24 h of food deprivation (Figures 1I and S1D). Therefore, QIH mice had lower blood glucose levels after 6 h of fasting and higher blood glucose levels after 24 h of fasting than control mice (Figures 1J and 1K). The same results were also observed in female mice, indicating that the effects of QIH on BW and blood glucose levels were not sexually dimorphic (Figures S3A and S3B).

**Figure 2.**
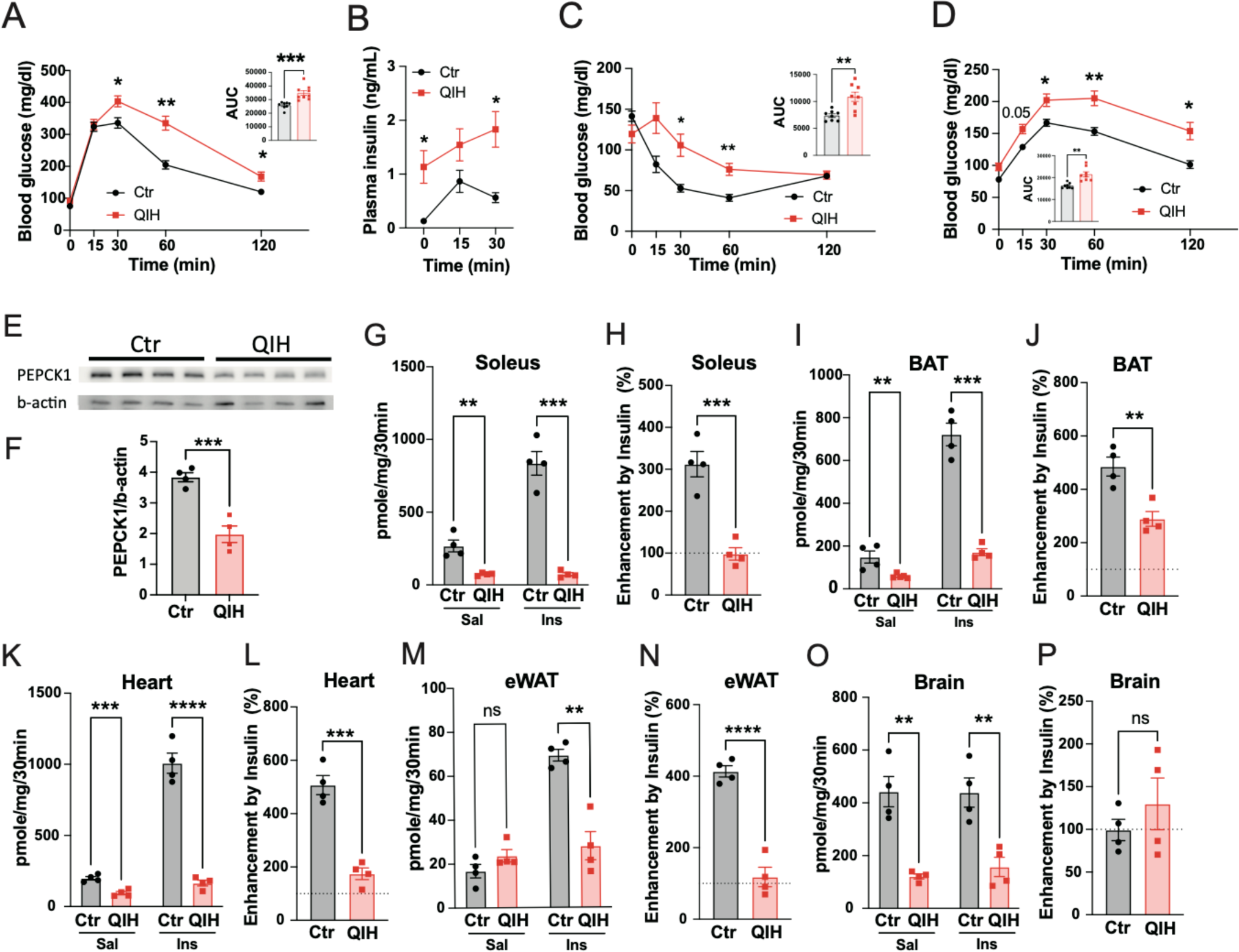
Glucose hypometabolic state in QIH mice. (A) Glucose tolerance test (GTT) of control (n = 8) and QIH (n = 9) mice (B) Plasma insulin levels in control (n = 6) and QIH (n = 6) mice during GTT (C) Insulin tolerance test (ITT) in control (n = 8) and QIH (n = 8) mice (D) Pyruvate tolerance test in control (n = 8) and QIH (n = 8) mice (E–F) Protein expression of liver PEPCK1 in control (n = 4) and QIH (n = 4) mice after 24-h fasting (E) Representative photos of immunoblotting (F) Quantification of liver PEPCK1 of control (n = 4) and QIH (n = 4) mice (G–P) 2DG uptake with or without insulin stimulation in the soleus (G–H), brown adipose tissue (BAT) (I–J), heart (K–L), epididymal white adipose tissue (eWAT) (M–N), and brain cortex (O–P) (G, I, K, M, O) Absolute 2DG uptake with saline or insulin injection in the soleus (G), BAT (I), heart (K), eWAT (M), and brain cortex (O) (H, J, L, N, P) Enhancement of 2DG uptake by insulin stimulation in the soleus (H), BAT (J), heart (L), eWAT (N), and brain cortex (P). All data are presented as means ± SEM; ns = not significant, *p < 0.05; **p < 0.01; ***p < 0.001; ****p < 0.0001.

Interestingly, plasma insulin levels were elevated after QIH induction and were unaffected by fasting, whereas plasma insulin levels in control mice decreased during fasting (Figure 1L). The higher insulin and blood glucose levels after 24 h of fasting strongly suggested that QIH mice were insulin resistant and glucose hypometabolic.

### QIH mice exhibit glucose hypometabolism

QIH mice showed hyperinsulinemia and higher fasting blood glucose levels, which are indicators of type II diabetes. We performed glucose and insulin tolerance tests (GTT and ITT) to investigate whether QIH mice developed glucose hypometabolism. The GTT indicated that QIH mice had lower glucose clearance ability and glucose intolerance (Figure 2A). Insulin levels during GTT were higher in QIH mice than those in control mice, indicating that impaired glucose tolerance was caused by insulin insensitivity rather than insufficient insulin levels (Figure 2B). Although insulin levels were high in QIH mice, exogenous glucose injection did not significantly increase insulin secretion (Figure 2B). As plasma insulin was abnormally high in QIH, we hypothesized that the central regulation of insulin secretion is disturbed. We examined c-fos expression in the paraventricular nucleus of hypothalamus (PVN) and observed that c-fos expression in the PVN particularly in oxytocin-expressing neurons was significantly decreased in QIH mice, suggesting that the suppressive signals of insulin secretion were decreased (Figure S4) (Papazoglou et al., 2022)

The ITT verified our hypothesis that QIH mice were insulin resistant (Figure 2C). Consistent with the GTT and ITT results, QIH mice were more pyruvate intolerant, increasing the possibility that QIH mice have higher gluconeogenesis after fasting (Figure 2D). However, the lower expression of liver PEPCK1 suggested lower gluconeogenesis in QIH mice (Figures 2E and 2F). We hypothesized that inhibited glucose utilization can compensate for the effects of decreased gluconeogenesis and result in pyruvate intolerance. Therefore, to evaluate glucose utilization in individual tissues, we used the 2DG uptake assay (Figures 2G–2P). Basal 2DG uptake was markedly decreased in the soleus, BAT, heart, and brain cortex of QIH mice (Figures 2G, 2I, 2K, and 2O). Interestingly, basal 2DG uptake in the eWAT was similar between the control and QIH groups (Figure 2M). Nevertheless, insulin-enhanced 2DG uptake was much lower in tissues from QIH mice, including eWAT (Figures 2G, 2I, 2K, and 2O). We further calculated the effects of insulin on 2DG uptake and observed that insulin effects were significantly lower in all tissues except the brain cortex between control and QIH mice (Figures 2H, 2J, 2L, 2N, and 2P). Notably, 2DG uptake was decreased in the brain cortex, which is insulin independent, suggesting that QIH not only affects insulin sensitivity but also disturbs insulin independent glucose metabolism.

Thus, QIH animals exhibited glucose hypometabolism with insulin resistance. Although hepatic gluconeogenesis was likely inhibited by QIH, the extremely low glucose uptake compensated for the decreased gluconeogenesis, thereby resulting in pyruvate intolerance.

### Body temperature abolishes QIH-mediated glucose homeostasis

As QIH mice exhibited hypothermia, glucose hypometabolism in QIH may be induced by hypothermia rather than by Qrfp^POA^-mediated circuits. Next, we investigated whether hypothermia is a factor for QIH-induced glucose hypometabolism by inducing QIH at 34°C to prevent a dramatic drop in body temperature (Figure 3A). At this temperature, QIH mice showed similar T_BAT_ to control mice after 24 h of fasting (Figures 3B and 3C). The efficacy of DREADD in mice after 24-h fasting at 34°C was confirmed by higher c-fos expression in Qrfp^POA:hM3Dq^ neurons (Figures 3D and 3E). To confirm that these activated Qrfp^POA^ remain functional to induce hypothermia, we transferred QIH mice from 34°C to 23°C to “cool down” these mice after 24-h fasting at 34°C. The T_BAT_ of QIH mice gradually decreased to 25°C, indicating that the activated Qrfp^POA^ remained able to induce QIH (Figures 3F and 3G).

**Figure 3.**
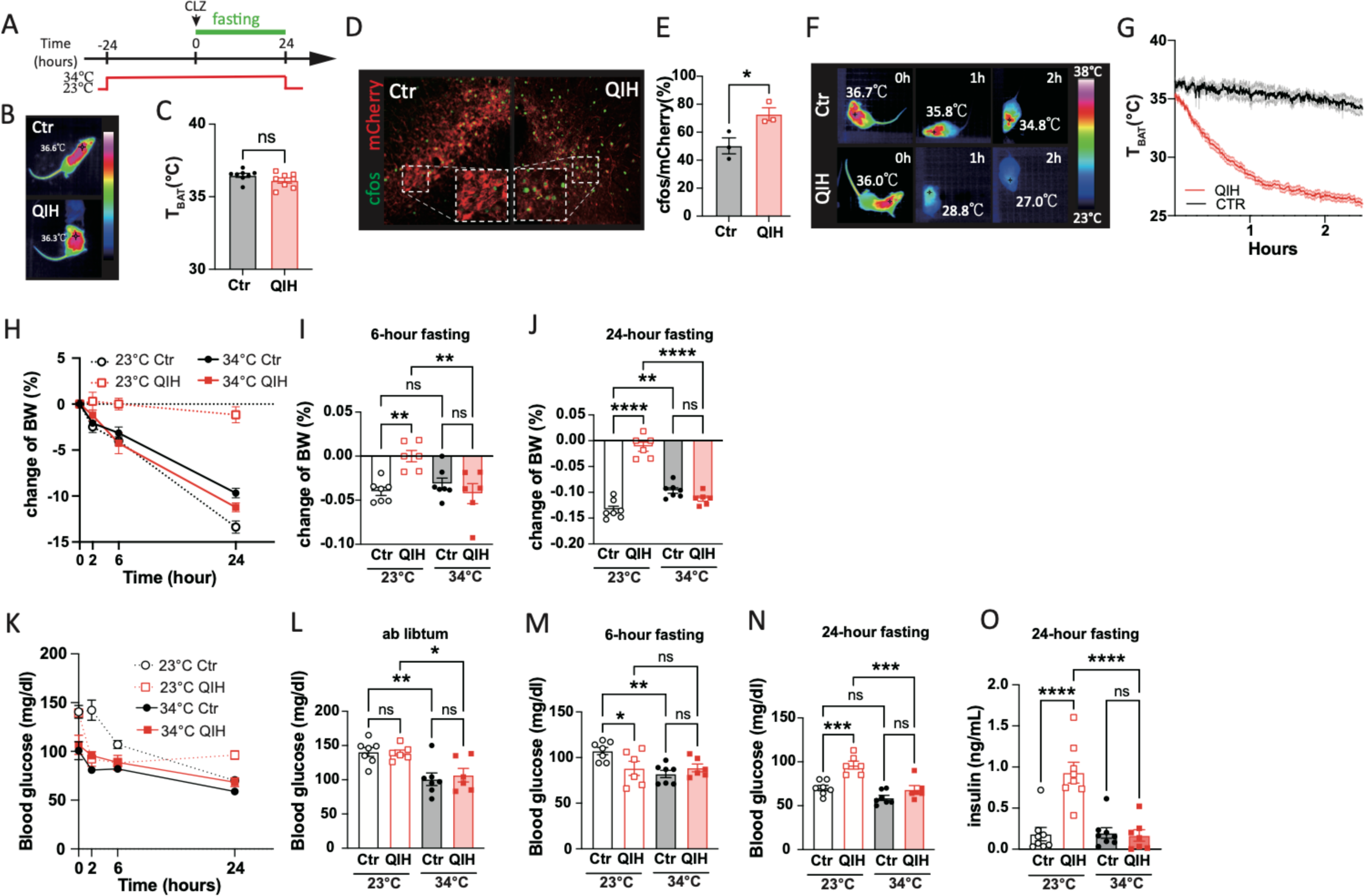
QIH-induced diabetes-like metabolic states are regulated by body temperature. (A) Diagram of the experimental design (B–C) Similar BAT temperature between control (n = 8) and QIH (n = 8) mice housed in 34°C after 24-h fasting (B) Representative thermograph of control and QIH mice (C) BAT temperature of QIH and control mice housed in 34°C (control, n = 8; QIH, n = 8) (D–E) cFos expression of Qrfp^POA^ in control (n = 3) and QIH (n = 3) mice housed after 24-h fasting in 34°C (D) Representative micrograph showing cFos expression in Qrfp^POA^ (E) Quantification of cFos expressing Qrfp^POA^ (F–G) T_BAT_ of control (n = 8) and QIH (n = 8) mice after being moved from 34°C to 23°C (F) Representative thermograph showing T_BAT_ of control and QIH mice (G) Traces of T_BAT_ in control and QIH mice (H–J) Changes of BW of control (n = 7) and QIH (n = 6) mice housed in 23°C or 34°C during fasting (H) Traces of BW changes in control and QIH mice (I–J) BW changes after (I) 6-h and (J) 24-h fasting (K–N) Blood glucose levels of control (n = 7) and QIH (n = 6) mice housed in 23°C and 34°C during fasting (K) Traces of blood glucose level changes in control (n = 7) and QIH (n = 6) mice (L–N) Blood glucose levels (L) before, (M) 6 h after, and (N) 24 h after QIH induction (O) Plasma insulin levels in control and QIH mice housed in 23°C (control, n = 8; QIH, n = 8) and 34°C (control, n = 8; QIH, n = 7) after 24-h fasting All data are presented as means ± SEM; ns = not significant, *p < 0.05; **p < 0.01; ***p < 0.001; ****p < 0.0001.

The control mice did not lose as much BW during fasting at 34°C as they did at 23°C, suggesting lower energy use at the warmer temperature (Figure 3H) (Clayton and McCurdy, 2018). However, QIH mice lost more BW at 34°C than that at 23°C, suggesting that energy utilization was higher at 34°C (Figures 3H–3J). Blood glucose levels were also measured during fasting after clozapine injection at 34°C to understand whether QIH-mediated antifasting glucose homeostasis still existed at higher temperatures (Figures 3K–3N). Surprisingly, blood glucose levels during fasting were similar between QIH and control mice at 34°C (Figures 3K–3N). Notably, basal blood glucose levels were lower in both groups after being housed at 34°C for 24 h before fasting and QIH induction (Figure 3L). In the control mice, blood glucose levels gradually decreased during fasting and were similar between housing at 23°C and 34°C after 24-h fasting, whereas blood glucose levels after 24-h fasting were significantly lower in QIH mice housed at 34°C than those housed at 23°C (Figure 3N). QIH mice had similar blood glucose levels to the control mice at 34°C during fasting, indicating that the QIH mice failed to reorganize glucose homeostasis at normal body temperature.

As high ambient temperature alters basal blood glucose levels (Figure 3L) (Clayton and McCurdy, 2018), prehousing animals at 34°C before QIH induction may alter the body situation and interfere with the effects of QIH on hypometabolism. To exclude this possibility, we housed mice at 23°C and transferred them to 34°C immediately after clozapine injection (Figure S5A). Blood glucose and BW between control and QIH mice were at the same level during fasting, indicating that acute elevation of body temperature is sufficient to abolish the hypometabolic effects of QIH (Figures S5B and S5C). Consistent with similar glucose levels, plasma insulin levels in 34°C-housed QIH mice were similar to those in control mice (Figure 3O).

### QIH-induced glucose hypometabolism is dependent on body temperature

Lower blood insulin levels and lower fasting blood glucose levels suggested that QIH mice had improved insulin sensitivity at 34°C. Therefore, we subsequently evaluated whether body temperature-mediated QIH-induced glucose hypometabolism. Glucose tolerance was not different between control mice housed at 23°C and 34°C, suggesting that ambient temperature does not significantly affect systemic glucose metabolism in normal mice (Dudele et al., 2015) (Figures 4A and 4B). However, the glucose tolerance of QIH mice was significantly improved at 34°C compared with that at 23°C and reached a level similar to that of the control groups (Figures 4A and 4B). The plasma insulin levels of QIH mice at 34°C did not differ from those of control mice during the GTT (Figure 4C). These results strongly indicated that the insulin sensitivity of QIH mice could be improved at 34°C. Consistent with this hypothesis, 34°C-housed QIH mice showed higher insulin sensitivity than 23°C-housed QIH mice with similar levels of 34°C-housed control mice (Figures 4D and 4E). Elevated ambient temperature did not change 2DG uptake in the soleus and eWAT in control mice, whereas it decreased 2DG uptake in the BAT and heart (Figures 4F–4I). Interestingly, increasing ambient temperature enhanced 2DG uptake by the brain cortex, whose metabolism is insulin independent (Figure 4J). However, 2DG uptake was higher in the soleus, eWAT, heart, and brain cortex of QIH mice at 34°C than at 23°C, suggesting that elevated body temperature increases glucose metabolism in QIH mice (Figures 4F, 4H, 4I, and 4J). The similar 2DG uptake levels between QIH and control mice validate the similar glucose metabolism observed in GTT and ITT (Figures 4A, 4B, 4D, and 4E). Notably, the heart, which is directly regulated by the POA, showed lower 2DG uptake than control mice at 34°C (Piñol et al., 2021) (Figure 4I). BAT 2DG uptake in QIH mice was not different between 23°C and 34°C, as BAT thermogenesis was inhibited by the warm environment (Figure 4G). Overall, the reversal of QIH-mediated glucose hypometabolism by elevated body temperature suggests that the effects of Qrfp^POA^ on glucose metabolism are secondary to hypothermia rather than Qrfp^POA^ regulation.

**Figure 4.**
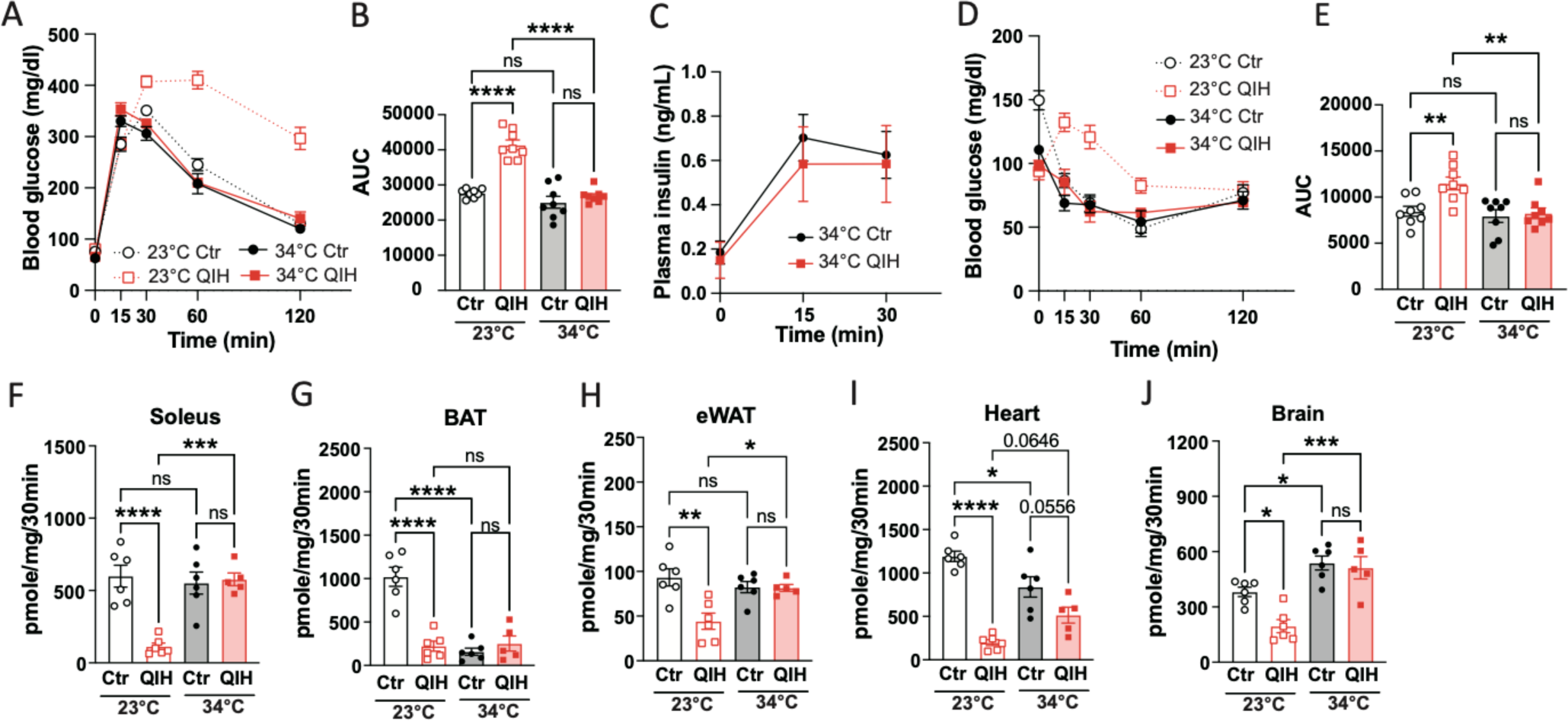
Increased body temperature abolishes QIH-mediated glucose hypometabolism. (A–B) GTT in control (n = 8) and QIH (n = 8) mice housed in 23°C and 34°C (A) Traces of blood glucose levels during GTT (B) Area under the curve (AUC) of (a) (C) Plasma insulin levels of control (n = 6) and QIH (n = 6) mice during GTT in 34°C (D–E) ITT in control (n = 8) and QIH (n = 8) mice housed in 23°C and 34°C (D) Trace of blood glucose levels during ITT (E) AUC of (D) (F–J) 2DG uptake in the (F) soleus, (G) BAT, (H) eWAT, (I) heart, and (J) brain cortex of control and QIH mice housed in 23°C (control, n = 6; QIH, n = 6) and 34°C (control, n = 6; QIH, n = 5) after insulin injection All data are presented as means ± SEM; ns = not significant, *p < 0.05; **p < 0.01; ***p < 0.001; ****p < 0.0001.

### Body temperature-mediated metabolic regulation is reversible

To exclude the possibility that higher ambient and body temperatures affect the turnover rate of clozapine within the body and affect the efficacy of QIH on glucose homeostasis, we cooled the body temperature of 34°C-housed QIH mice from euthermia to hypothermia by transferring them to room temperature (23°C) after fasting. Mice housed at 34°C were injected with clozapine and deprived of food. These mice were transferred to 23°C for cooling after 4-or 16-h fasting (Figures 5A and 5G). Once the mice were exposed to 23°C, T_BAT_ decreased and could be as low as approximately 25°C within 2 h (Figures 5B and 5H). Blood glucose levels after 6-h fasting (4 h at 34°C followed by 2 h at 23°C) were increased in control mice after exposure at 23°C but were not changed in QIH mice (Figure 5C). This increase resulted in higher blood glucose levels in cooled control mice than in cooled QIH mice. Interestingly, among 18-h fasting mice, QIH mice developed increased blood glucose levels after 2 h of cooling; however, no differences were observed in control mice with or without cooling (Figure 5I). Therefore, the blood glucose levels of the cooled QIH mice were higher than those of the cooled control mice after overnight fasting. These results of blood glucose levels in cooled mice were similar to those of 23°C-housed control and 23°C-housed QIH mice (Figure 1H). Therefore, the rapid changes in blood glucose levels suggest that glucose homeostasis is tightly and promptly regulated by either ambient or body temperatures (Figures 3K–3N). The glucose tolerance of control mice was not affected by cooling and was at the same level as that of 34°C-housed QIH mice (Figures 5J and 5K). Nevertheless, QIH mice were significantly more glucose intolerant after cooling than those in the other groups (Figures 5J and 5K). Consistent with these results, the insulin sensitivity of QIH mice was remarkably reduced after 2-h cooling, whereas control mice showed similar insulin sensitivity (Figures 5D and 5E). Although 6-h–fasted control mice had higher basal blood glucose levels after cooling (Figures 5C and 5D), minimal blood glucose levels during the ITT were similar between control mice with and without cooling, indicating that insulin sensitivity was not altered by ambient temperature (Figure 5F). Again, the higher minimal blood glucose levels of cooled QIH mice during the ITT indicated insulin resistance in these mice (Figure 5F). Taken together, body temperature modulation can efficiently regulate glucose metabolism. Although changes in ambient temperature seem to alter glucose homeostasis, glucose tolerance and insulin sensitivity could not be remodeled without large changes in body temperature.

**Figure 5.**
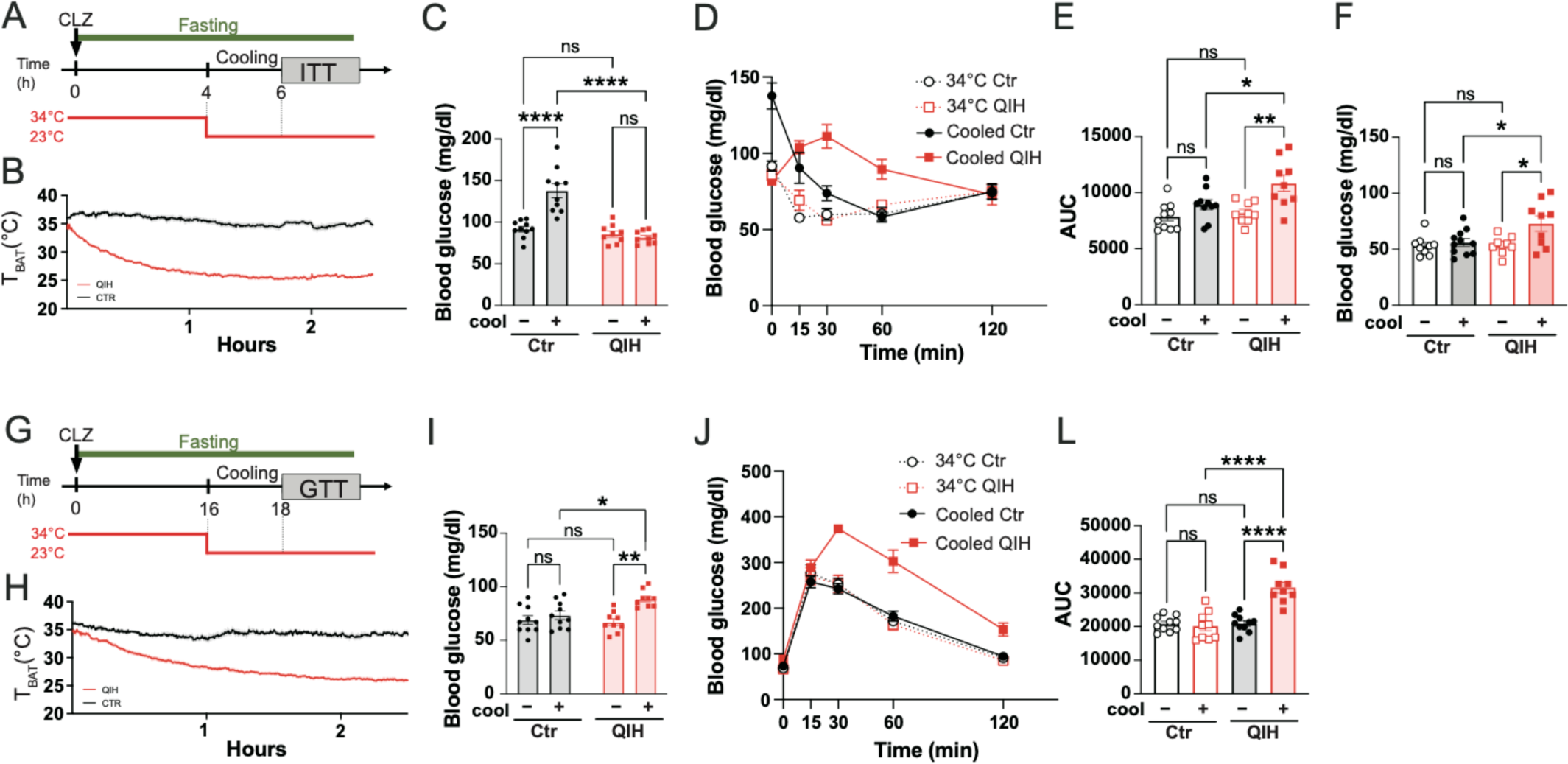
Acute cooling down of body temperature recovers QIH-mediated glucose hypometabolism. (A, G) Diagram of experimental design for 6-h (A) and 18-h (G) fasting (B) Traces of body temperature of 6-h–fasted control (n = 8) and QIH (n = 8) mice after being moved to 23°C (C) Blood glucose levels of 6-h–fasted control (n = 10) and QIH (n = 9) mice before and after 2-h cooling (D–E) ITT in 34°C-housed control (n = 10) and QIH (n = 9) mice with or without cooling (D) Traces of blood glucose levels during ITT (E) AUC of (D) (F) Minimal glucose level during ITT (H) Traces of body temperature of 18-h–fasted control (n = 8) and QIH (n = 8) mice after being moved to 23°C (I) Blood glucose levels of 18-h–fasted control (n = 10) and QIH (n = 9) before and after 2-h cooling (J–K) GTT in 34°C-housed control (n = 10) and QIH (n = 9) mice with or without cooling (J) Traces of blood glucose levels during GTT (K) AUC of (I) All data are presented as means ± SEM; ns = not significant, *p < 0.05; **p < 0.01; ***p < 0.001; ****p < 0.0001.

### Body temperature is a critical factor in tuning feeding and locomotion behaviors

QIH mice are extremely inactive and inappetent at room temperature (Takahashi et al., 2020) (Figure 1F). However, to compensate for the increased energy utilization of QIH mice at 34°C, energy intake should be increased (Figures 3 and 4). Therefore, we investigated the effect of body temperature on food intake, which was strongly inhibited in room temperature-housed QIH mice. Consistent with previous studies, the 24-h food intake of control mice was reduced as ambient temperature increased (Figure 6A) (Cui et al., 2016; John et al., 2022). QIH mice, which were inappetent at 23°C, markedly increased food intake at 34°C although remained lower than that of 34°C-housed control mice (Figure 6A). Refed food intake showed a pattern similar to that of 24-h food intake (Figures 6B and 6C). In control mice, refed food intake after 24-h fasting was higher at 23°C than that at 34°C (Figure 6C). The appetite of QIH mice also recovered at 34°C; however, the cumulative refed food intake remained lower than that of 34°C-housed control mice (Figure 6C).

**Figure 6.**
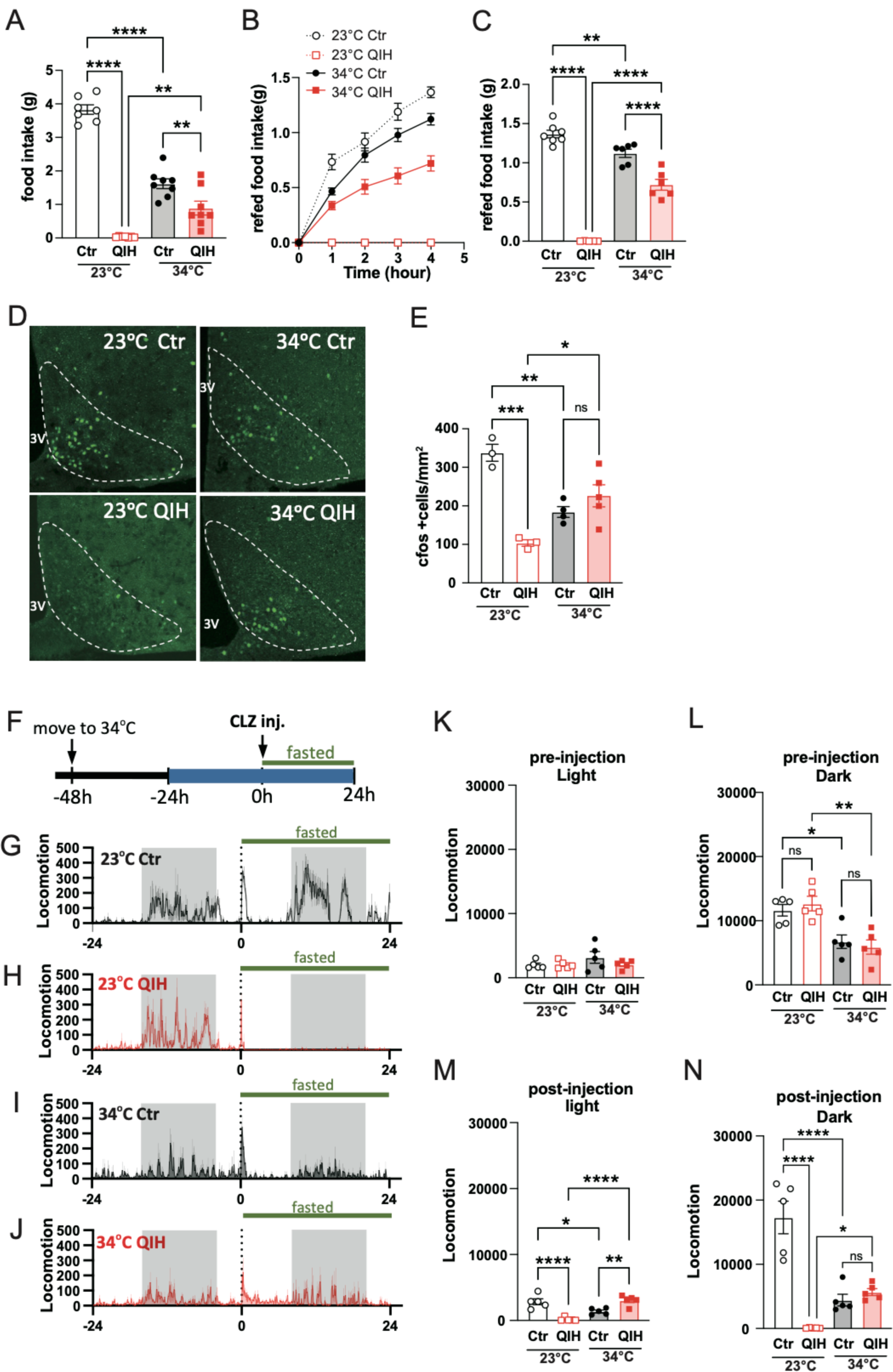
Body temperature regulates food intake and locomotion. (A) Twenty-four-hour food intake of control and QIH mice housed in 23°C (control, n = 7; QIH, n = 7) and 34°C (control, n = 8; QIH, n = 8) (B–C) Accumulative refed food intake of control and QIH mice housed in 23°C (control, n = 7; QIH, n = 7) and 33°C (control, n = 6; QIH, n = 6) after 24-h fasting (B) Traces of accumulative refed food intake (C) Food intake after 4-h refed (D–E) Immunostaining of cFos in the arcuate nucleus (ARC) of control and QIH mice housed in 23°C (control, n = 3; QIH, n = 3) and 34°C (control, n = 4; QIH, n = 4) (F–N) Locomotion activity of control (n = 5) and QIH (n = 5) mice housed in 23°C and 34°C (F) Experimental design to measure the locomotion activity of control and QIH mice. The blue bar indicates the recording period. (G–I) Traces of locomotion activity of control and QIH mice housed in 23°C (G–H) and 33°C (I–J) from 24 h before to 24 h after food deprivation (K–L) Locomotion activity during (K) light and (L) dark periods before food deprivation (M–N) Locomotion activity during (M) light and (N) dark periods after food deprivation and QIH induction All data are presented as means ± SEM; ns = not significant, *p < 0.05; **p < 0.01; ***p < 0.001; ****p < 0.0001.

The arcuate nucleus (ARC) is the core region of food intake regulation in which proopiomelanocortin neurons and agouti-related peptide (Agrp) neurons inhibit or enhance food intake, respectively (Deem et al., 2022; Vicent et al., 2018). We investigated whether ambient or body temperature alters neuronal activity in the ARC. After 24-h fasting, control mice showed abundant c-fos expression in the ARC at 23°C, particularly in the medial ARC where Agrp neurons aggregate, reflecting the hunger sensation induced by 24-h fasting (Figures 6D and 6E). However, fasted QIH mice had significantly lower ARC c-fos expression than control mice, which is consistent with the anorexic effects of QIH at 23°C (Figures 6D and 6E). When mice were housed at 34°C, control mice showed decreased c-fos expression in the ARC, suggesting milder hunger stimulation at higher ambient temperature, which is consistent with the results of reduced food intake at 34°C (Figure 6A). The c-fos expression in the ARC of QIH mice was significantly increased at 34°C, supporting the result that QIH mice had better appetite at higher body temperatures (Figures 6D and 5E).

Subsequently, we hypothesized that QIH mice should be more physically active owing to higher food intake (Figures 6B–6D), higher glucose metabolism, and greater BW loss at 34°C (Figures 3J and 4). We implanted the nanotag^®^ probe, a body-implantable actimeter, in the abdomen for monitoring locomotor activity housed at 23°C or 34°C before and after QIH induction (Figures 6F–6J). Before fasting and clozapine injection, either QIH or control mice developed decreased locomotor activity at 34°C compared with that at 23°C during the dark but not the light period (Figures 6K and 6L) (Zahm et al., 2014). After clozapine injection, as expected, the locomotor activity of QIH mice was significantly lower than that of control mice at 23°C in both light and dark periods (Figures 6M and 6N). The locomotor activity of QIH mice after clozapine injection was significantly higher at 34°C than that at 23°C, indicating that body temperature is a factor in regulating locomotor activity (Figures 6M and 6N). Interestingly, QIH mice showed higher locomotor activity than control mice after clozapine injection at 34°C, suggesting that Qrfp^POA^ activation directly enhances locomotor activity via a temperature-independent mechanism (Figures 6M and 6N).

In conclusion, we noted that QIH reorganizes glucose homeostasis with a stable but extremely insulin resistant metabolic status. However, this diabetes-like status is regulated by body temperature and not by neuronal circuits involving Qrfp^POA^. Therefore, Qrfp^POA^ plays a role in lowering body temperature, and hypometabolism and torpid behavior are secondary to hypothermia (Figure 7).

**Figure 7.**
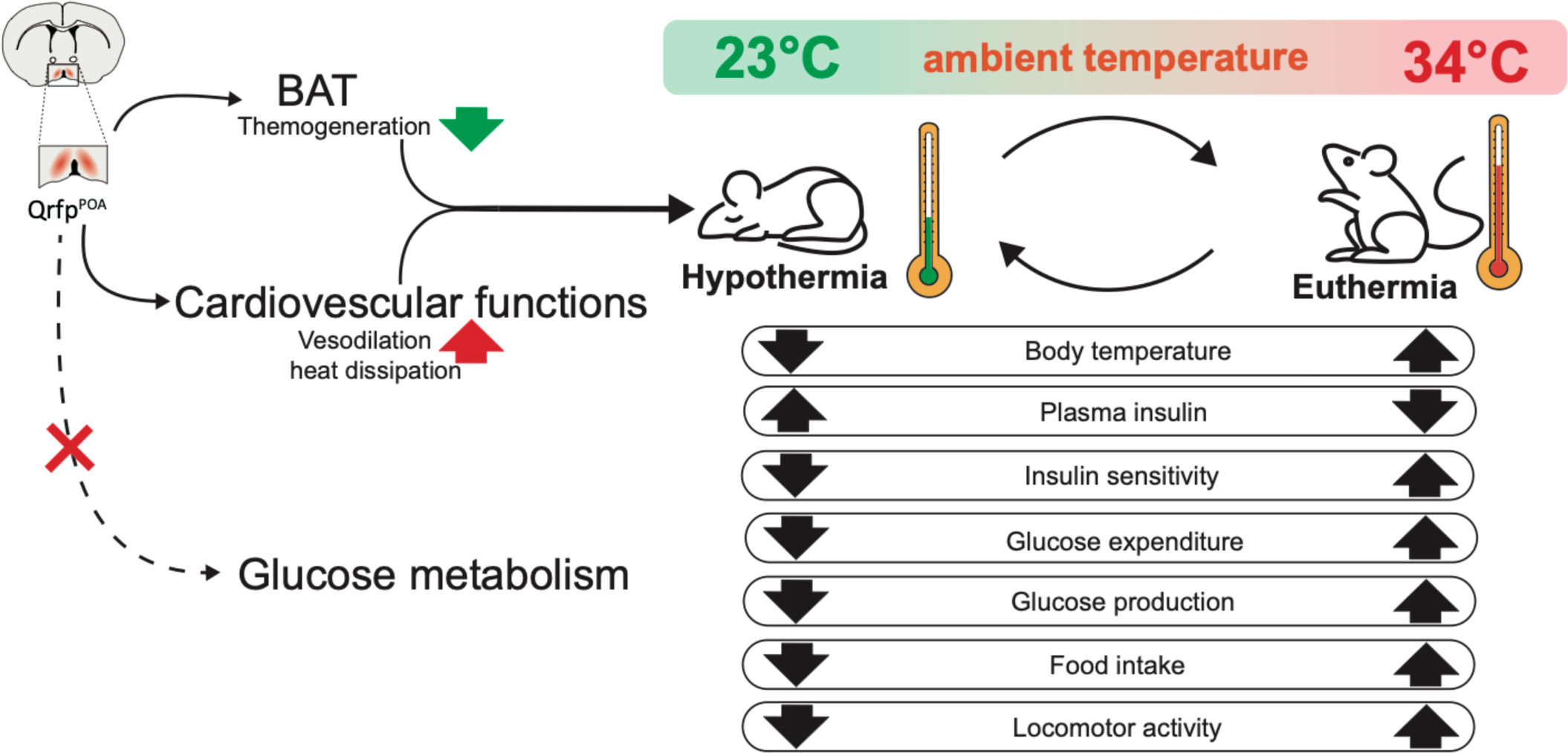
Hypothermia regulates glucose metabolism. Qrfp^POA^ activation induces hypothermia by decreasing BAT thermogenesis and increasing vasodilation without directly affecting glucose metabolism. Hypothermia results in glucose hypometabolism with increasing plasma insulin and decrease insulin sensitivity. Hypothermic mice show inappetance and low locomotor activity. The QIH mediated glucose hypometabolism and behaviors are abolished by increasing body temperature to euthermia.

## Discussion

QIH is a state of hypothermia and hypometabolism caused by Qrfp neuron activation in the POA. From QIH onset, blood glucose levels decreased along with the decrease in body temperature and were maintained at stable levels. Surprisingly, despite the decrease in blood glucose levels, plasma insulin increased to abnormally high levels as soon as the animal entered the QIH status. This is consistent with the previous notion that QIH is a hibernation-like status rather than torpor because hyperinsulinemia was also observed in mammalian hibernators but not in torpid mice (Buck et al., 2002; Florant et al., 1985). Moreover, the stable and fasting-disregard blood glucose levels and hyperinsulinemia suggest that the reorganized glucose homeostasis in QIH is not responsive to body energy states. The brain particularly hypothalamic nuclei such as the ARC, ventromedial hypothalamus (VMH), dorsomedial hypothalamus (DMH), and PVN play a critical role in sensing blood glucose, which can reflect energy levels to regulate whole-body energy metabolism (Alvarsson and Stanley, 2018; Huang et al., 2022; Shimazu and Minokoshi, 2017). We showed that QIH universally silences neuronal activity, at least in the ARC. In fact, we also observed that neuronal activity in the VMH, DMH, and PVN is significantly inhibited during QIH. These nuclei may lose their functions to detect energy levels, thereby leaving stable blood glucose and insulin levels during fasting. Moreover, blood insulin levels are regulated by the hypothalamus (Heni et al., 2020; Pozo and Claret, 2018). Hypothalamic neuron inhibition, at least oxytocin neurons in the PVN, may lead to hyperinsulinemia in QIH mice (Papazoglou et al., 2022). Decreased insulin degradation is unlikely to be the cause of hyperinsulinemia in QIH mice because exogenous glucose injection did not adequately increase insulin levels. Thus, insulin secretion dysregulation may be the main reason for abnormally high plasma insulin levels. Pancreatic islets function during QIH/hypothermia will be a significant issue in future studies.

Blood glucose levels are determined by plasma insulin levels and insulin sensitivity. As expected, QIH mice showed glucose hypometabolism with insulin resistance at room temperature because both blood glucose and insulin levels were simultaneously elevated. An interesting question is how the new glucose homeostasis develops in QIH. The rapid decrease in blood glucose levels during early QIH, when hyperinsulinemia develops, suggests that mice can have normal insulin responsiveness at the very beginning of QIH. Therefore, it is more reasonable that the increase in insulin levels precedes insulin resistance development in QIH animals; however, hibernation hyperinsulinemia is believed to be a consequence of insulin resistance (Wu et al., 2013). Reduced insulin sensitivity in QIH mice has evolved to compensate for the effects of hyperinsulinemia. In addition, insulin stimulates nitric oxide production in endothelial cells, thereby resulting in vasodilation that may accelerate the decrease in body temperature (Almeida and Branco, 2001; Wu and Meininger, 2009). Thus, hyperinsulinemia in early QIH may contribute to vasodilation and lower body temperature. Therefore, insulin responsiveness between different stages of QIH may be a significant issue for future studies.

Metabolism, such as oxygen consumption and energy use, is affected by ambient temperature (Seeley and MacDougald, 2021; Zhao et al., 2022). However, despite oxygen consumption decreases at warm temperature, changing the ambient temperature is insufficient to alter glucose tolerance and fasting insulin levels in lean mice (Clayton and McCurdy, 2018; Dudele et al., 2015). Nevertheless, glucose hypometabolism in QIH mice was abolished by warming these animals to normal body temperature. These results suggest that glucose hypometabolism was mainly caused by hypothermia rather than regulation by Qrfp^POA^. Therefore, Qrfp^POA^ does not affect glucose metabolism; instead, these neurons decrease body temperature, which affects glucose metabolism. In support of this, in a rat model, elevating muscle temperature increased glucose utilization, whereas hypothermia decreased glucose utilization (Fuhrman and Fuhrman, 1963; Koshinaka et al., 2013).

Among the tissues used to measure 2DG uptake, only heart 2DG uptake was lower than that in the control at 34°C, indicating that Qrfp^POA^ directly control cardiac function rather than other tissues. Consistent with this, different populations of neurons in the POA increase or decrease heart rate and cardiovascular function (Duarte et al., 2017; Hunt et al., 2010; Piñol et al., 2021). Although QIH mice had a markedly low heart rate, which was also noted in hypothermia induced by estrogen receptor-expressing neurons in the POA (Zhang et al., 2020), whether bradycardia is controlled by these neurons or by hypothermia needs further studies. Besides body temperature, the POA regulates locomotor activity (Brudzynski and Mogenson, 1986; Ogawa et al., 2003; Xu et al., 2023; Zhang et al., 2023). The locomotor activity of QIH mice, which was extremely inhibited at 23°C, was higher than that of the control at 34°C, suggesting that Qrfp^POA^ activation enhanced locomotion, and this enhancement was masked by hypothermia in QIH mice (Sinnamon, 1992, 1993). Therefore, hypothermia may interfere with phenotypes of POA-regulated behaviors or physiology and complicate the interpretation of results. In recent decades, the POA has been reported to regulate physiology and behaviors, including sleep, feeding, and parental behavior (Kohl and Dulac, 2018; Rothhaas and Chung, 2021). Disturbed body temperature can sometimes be induced by manipulating POA neurons and interferes with the actual functions of the POA. The extent to which POA-mediated hypothermia or hyperthermia disrupts these physiology and behaviors may be a question for future studies.

## Acknowledgment

We are grateful to the members of the Biophotonics Laboratory and the Transformative Research Area ‘‘Hibernation Biology” for helpful discussions on this study. We also thank Yuki Watakabe, Miwa Kawachi, Maki Watanabe, Miyoko Shimomura, and Chiemi Hyodo for their help with animal care and laboratory management, and Kana Tsuchiya and Ayaka Osamura for their excellent secretary work. We also appreciate Prof. Takeshi Sakurai (University of Tsukuba) and Genshiro Sunagawa (RIKEN BDR) for sharing Qrfp-iCre mice with us, and Prof. Makoto Tominaga (ExCELLS/NIPS) and Prof. Yoshifumi Yamaguchi (Hokkaido University) for the valuable discussion.

## Author contributions

M.L. and R.E. designed the research project. M.L. and C.C. performed the experiments. M.L. performed the data analysis. C.T., T.N., and R.E. supervised the research. All authors contributed to the writing of the manuscript and approved the submitted version.

## RESOURCE AVAILABILITY

### Lead contact

Further information and requests for resources and reagents should be directed by the lead contact Ryosuke Enoki

### Materials availability

This study did not generate new unique reagents

### Data and code availability

All data needed to evaluate the conclusions in the study are presented in the main text and Supplementary Materials. Further study-related data can be requested from the authors.

## EXPERIMENTAL MODEL AND SUBJECT DETAILS

### Mouse model

Qrfp-iCre mice were transported from the RIKEN BioResource Research Center (BRC No. RBRC11137). Animals were housed at 22°C–24°C and 30%–60% humidity with a 12-h light/12-h dark cycle and provided food and water access *ad libitum.* Eight-to 10-week-old male Qrfp-iCre mice were performed viral injection and followed experiments. All animal studies were performed on the basis of the ARRIVE guidelines; all animal care and experimental procedures were approved by the Institutional Animal Care and Use Committee of the National Institute of Natural Sciences and were performed according to the guidelines of the National Institute for Physiological Science (Approval 23A062).

## METHOD DETAILS

### Surgery for virus injection and nanotag^®^ implantation

Eight-to 10-week-old male Qrfp-iCre mice were placed on the stereotaxic instrument after anesthesia with a ketamine (100 mg/kg) and xylazine (10 mg/kg) mixture. Adeno-associated virus (AAV) (AAV5-DIO-syn-hM3Dq-mCherry or AAV5-DIO-syn-mCherry, addgene) was bilaterally injected into the POA with 300 nL each side in the coordinates of the anterior–posterior (AP) direction, 0.38; medial–lateral (ML), ± 0.3; and dorsal–ventral (DV), − 5.25. The titer of all AAV was over 7.0 × 10^12^ genome copies/mL. The open wound was sutured following viral injection. Injection efficacy was tested by intraperitoneal injection of clozapine (0.2 mg/kg; *MilliporeSigma, Burlington, MA, USA*) 4 weeks postoperatively. The mice were excluded when the BAT temperature was higher than 30°C after 24 h of clozapine injection. To measure the locomotion activity, a nanotag^®^ (Kissei Comtec Ltd., Nagano, Japan) was implanted into the abdominal cavity. After anesthesia with the ketamine (100 mg/kg) and xylazine (10 mg/kg) mixture, the abdominal cavity was opened by an incision at the middle line. The nanotag^®^ was placed inside the abdominal cavity, and the incision was closed. Mice were recovered at least 1 week before the experiments.

### Recording of the temperature

To record the BAT temperature (T_BAT_), an infrared FLIR AX5 thermal-imaging camera (Teledyne FLIR, Wilsonville, OR, USA) was positioned 40 cm above experimental cages. The back hair was shaved on the day before recording the temperature. To record changes in BAT temperature under QIH, mice were injected with clozapine (0.2 mg/kg) and immediately moved into a recording chamber under an infrared thermal-imaging camera. Mice were provided with water and food in the recording cage unless otherwise stated. Thermograms were collected at 0.1 Hz, and the highest temperature in the chamber was defined as T_BAT_. To record the T_BAT_ of mice during the cooling period, the mice were placed under a thermal-imaging camera immediately after moving from the 34°C chamber (HC-10, ShinFactory, Fukuoka, Japan) to 23°C.

### Body weight (BW), tissue weight, and blood glucose levels during fasting

Mice were injected with clozapine (0.2 mg/kg) at 10:00 a.m. and moved into clean cages without food. BW and blood glucose levels were measured at 0, 2, 6, 18, and 24 h after food deprivation and clozapine injection. To measure blood glucose levels, a small incision was made on the tail tip. A drop (approximately 2 μL) of blood from the incision was used to measure blood glucose levels using a handheld glucometer (Nipro Free style, Nipro, Osaka, Japan). To measure tissue weight, mice were sacrificed 24 h after clozapine injection and fasting. Tissues including those in the soleus, gastrocnemius, interscapular brown adipose tissue (BAT), epididymal white adipose tissue (eWAT), inguinal WAT, mesenteric WAT, liver, stomach, and intestine were carefully collected and weighed.

### Glucose, insulin, and pyruvate tolerance tests

For the glucose tolerance and pyruvate tolerance tests, mice were injected with clozapine and fasted for 18 h. Each animal received 2 g/kg BW of glucose for the glucose tolerance test (GTT) and 2 g/kg BW of sodium pyruvate for the pyruvate tolerance test by intraperitoneal injection. For the insulin tolerance test, mice were injected with clozapine and fasted for 6 h. Each animal received 0.5 U/kg BW of insulin by intraperitoneal injection. Blood glucose levels were measured at 0, 15, 30, 60, and 120 min after injection.

### Measurement of plasma insulin levels

To measure plasma insulin during QIH, 50 μL of blood was collected from the tails at 0, 6, and 24 h after clozapine injection. To assess glucose-stimulated insulin secretion, glucose (2 g/kg) was intraperitoneally injected, and blood samples (50 μL) from the tail were collected at 0, 15, and 30 min after glucose injection. Blood samples were mixed with 2 μL of heparin (200 U/mL). Plasma was collected after centrifugation for 10 min at 1,000 × g and maintained at −80°C until insulin was measured. Insulin concentration was measured using a Mouse Insulin ELISA KIT (FUJIFILM Wako, Osaka, Japan), and all procedures were performed following the protocols provided in the kit.

### Immunoblotting

Mice were sacrificed after clozapine injection and fasting for 24 h. A small piece of the liver was collected and immediately frozen at −80°C until experiments. Tissues were homogenized in RIPA buffer at 4°C by sonication. After centrifugation, the supernatant was fractionated by SDS-PAGE, and proteins were transferred to a polyvinylidene fluoride membrane. After blocking with 5% skimmed milk, the membrane was incubated with 1 μg/mL rabbit anti-PEPCK (Abcam, Cambridge, UK) or mouse anti-beta-actin (Cell Signaling Technology, Danvers, MA, USA) at 4°C overnight. Immune complexes were visualized using horseradish peroxidase-linked goat anti-rabbit immunoglobulin and enhanced chemiluminescence reagents (GE Healthcare, Tokyo, Japan).

### Measurement of 2DG uptake

To assess glucose utilization of tissues, 2DG uptake was measured. Mice were injected with clozapine to induce QIH and fasted for 18 h. Each animal was injected with 5 μmole of 2DG 10 min after insulin (0.75 U/kg BW) or saline injection. Mice were sacrificed 30 min after 2DG injection, and tissues such as those in the soleus, BAT, eWAT, heart, and brain cortex were collected and stored at −80°C until experiments. Tissues were homogenized in 500 μL of 10 mM Tris-HCl (pH, 8.0) by sonication. Samples were subsequently incubated at 95°C for 15 min, followed by centrifugation for 10 min at 16,000 × g. The supernatant was collected to measure 2DG uptake using the 2DG Uptake Measurement Kit (Cosmo Bio, Japan).

### Immunohistochemistry

Mice at 23°C or 34°C were injected with clozapine to induce QIH and were moved to clean cages without food. After 24-h fasting, mice were perfused with 0.1-M phosphate buffer (PB) followed by 4% paraformaldehyde transcardially. Brain sections (50 μm each) containing the POA were collected. The floating sections were incubated with rabbit anti-cFos antibody (1:2,000; Cell Signaling Technology, Danvers, MA, USA) or mouse anti-mCherry antibody (1:1,000; Abcam, Cambridge, UK) in staining solution (0.1-M PB containing 4% normal goat serum, 0.1% glycine, and 0.2% Triton X-100) overnight at room temperature (23°C). After rinsing with PB, the sections were incubated in secondary antibody (1:500, Alexa 488 or 594 Goat Antirabbit; Cell Signaling Technologies, Danvers, MA, USA) for 2 h at room temperature. Stained sections were washed with PB three times and mounted on glass slides with Vectashield (Vector Laboratories, Burlingame, CA, USA).

### Recording of locomotor activity

Locomotion activity was recorded using a nanotag^®^ (Kissei Comtec Ltd., Nagano, Japan). Mice were housed at 34°C or 23°C for 1 day before recording for acclimatization. A total of 48 h of locomotion activity was recorded for each experiment. Recording was started 24 h before clozapine injection and fasting (-24 h) and was continued until 24 h after clozapine injection (24 h). At the time of clozapine injection (0 h), mice received intraperitoneal injection of 0.2-mg/kg clozapine and were moved to clean cages without food. Data from the first hour after injection were excluded because hyperactivity was induced by injection.

## QUANTIFICATION AND STATISTICAL ANALYSIS

To determine the effect of QIH and body/ambient temperature, two-way or one-way analysis of variance (ANOVA) was used using Prism 10 software (GraphPad). For repeated-measures analysis, ANOVA was used when values over different times were analyzed, followed by Sidak multiple comparison tests. When only two groups were analyzed, statistical significance was determined using unpaired Student’s *t*-test (two-tailed *p* value). A *p* value of <0.05 was considered statistically significant. Quantification of c-fos positivity in PVN and ARC were done in blinded fashion. All data were presented as means ± SEM. The representative image was obtained from at least three independent experiments.

---

**Supplementary Figure 1.**
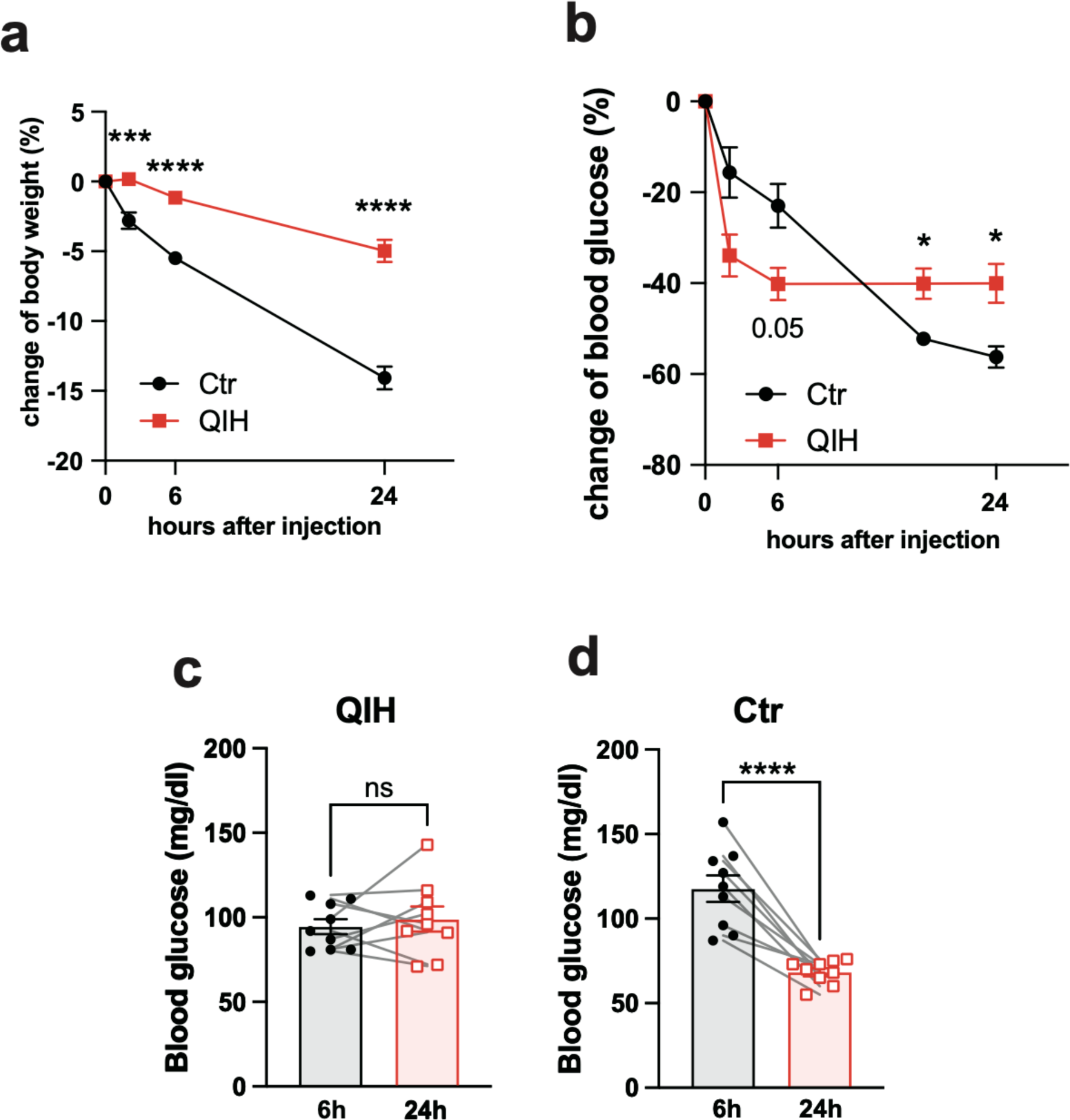

**Supplementary Figure 2.**
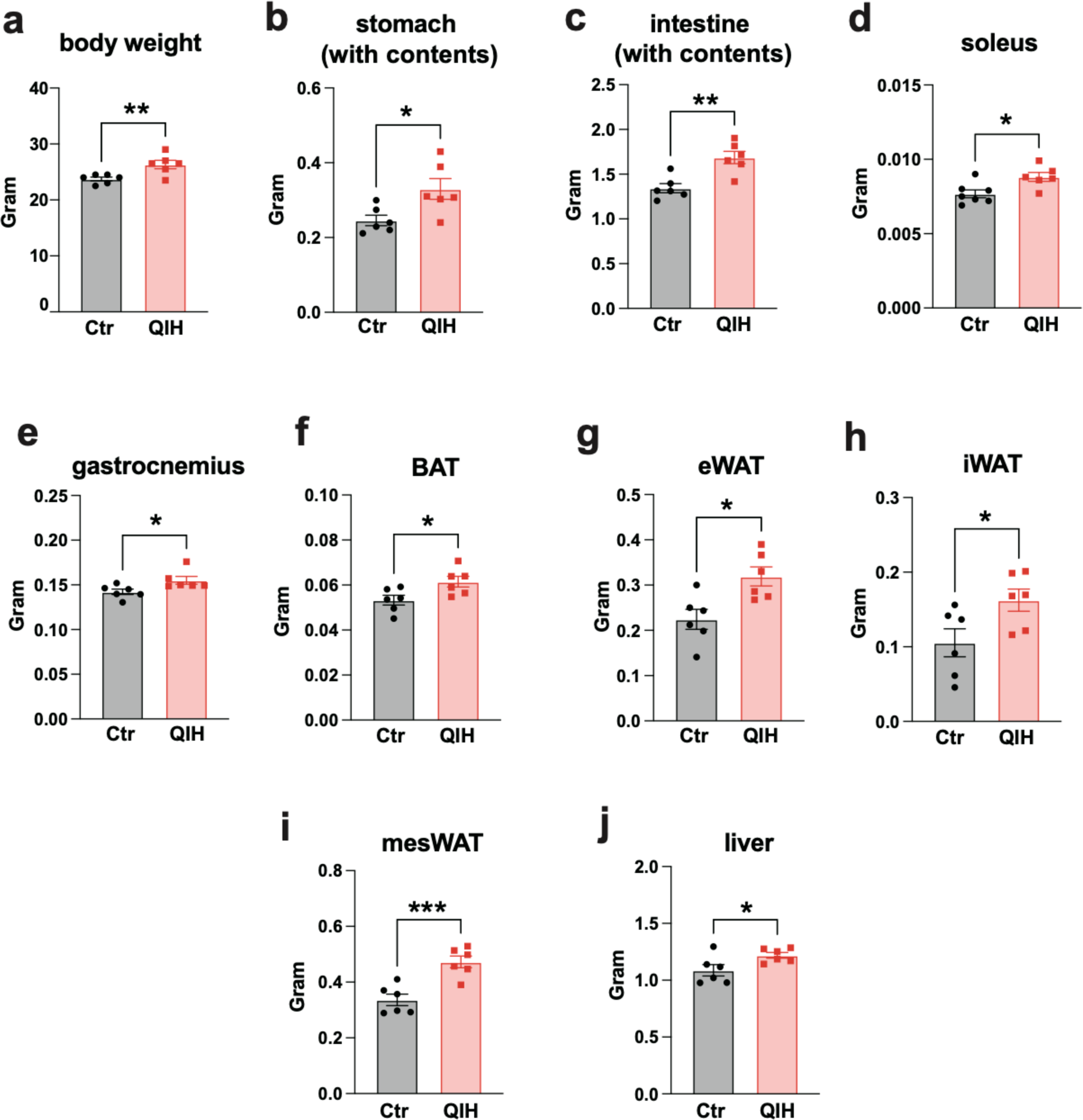

**Supplementary Figure 3.**
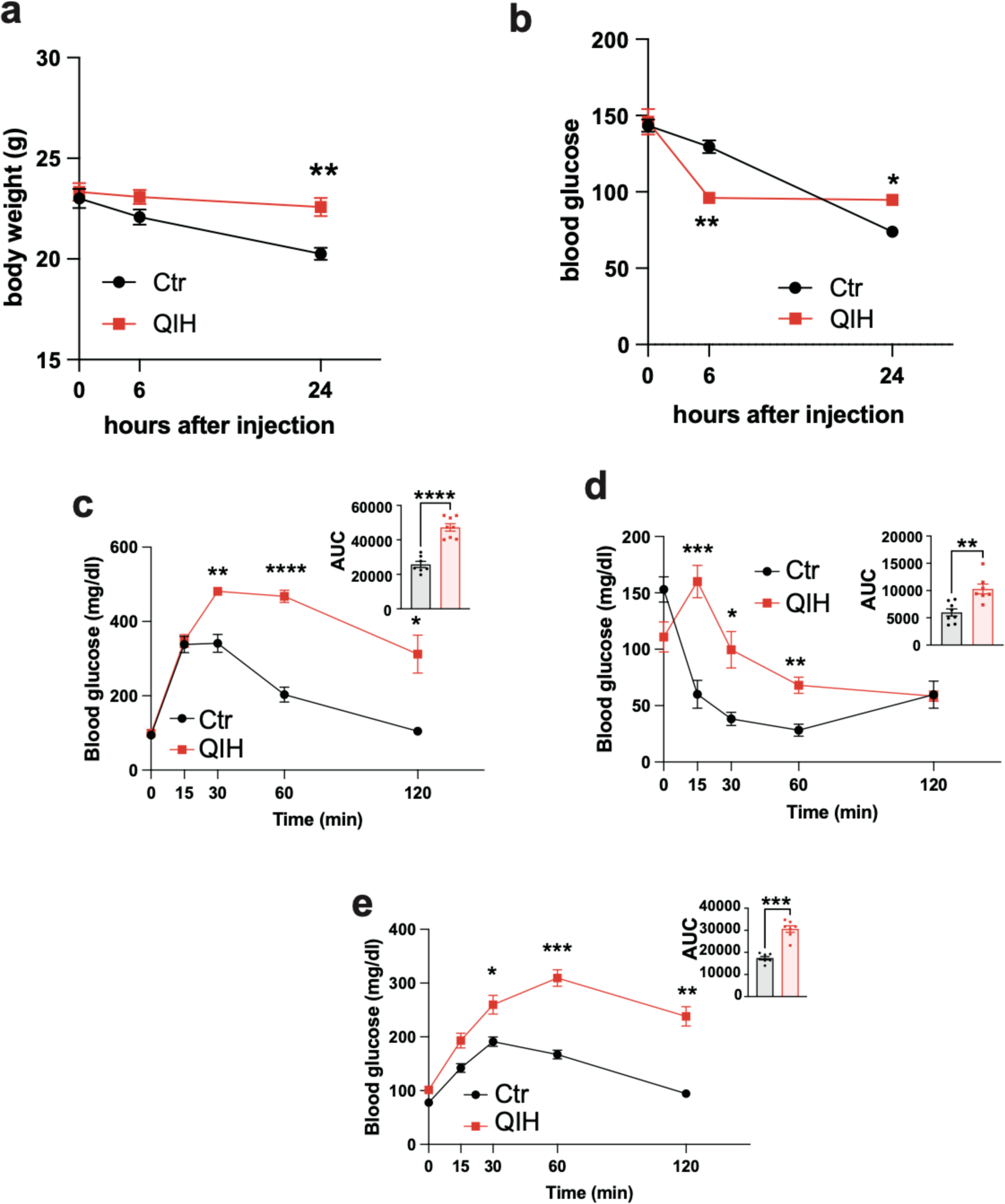

**Supplementary Figure 4.**
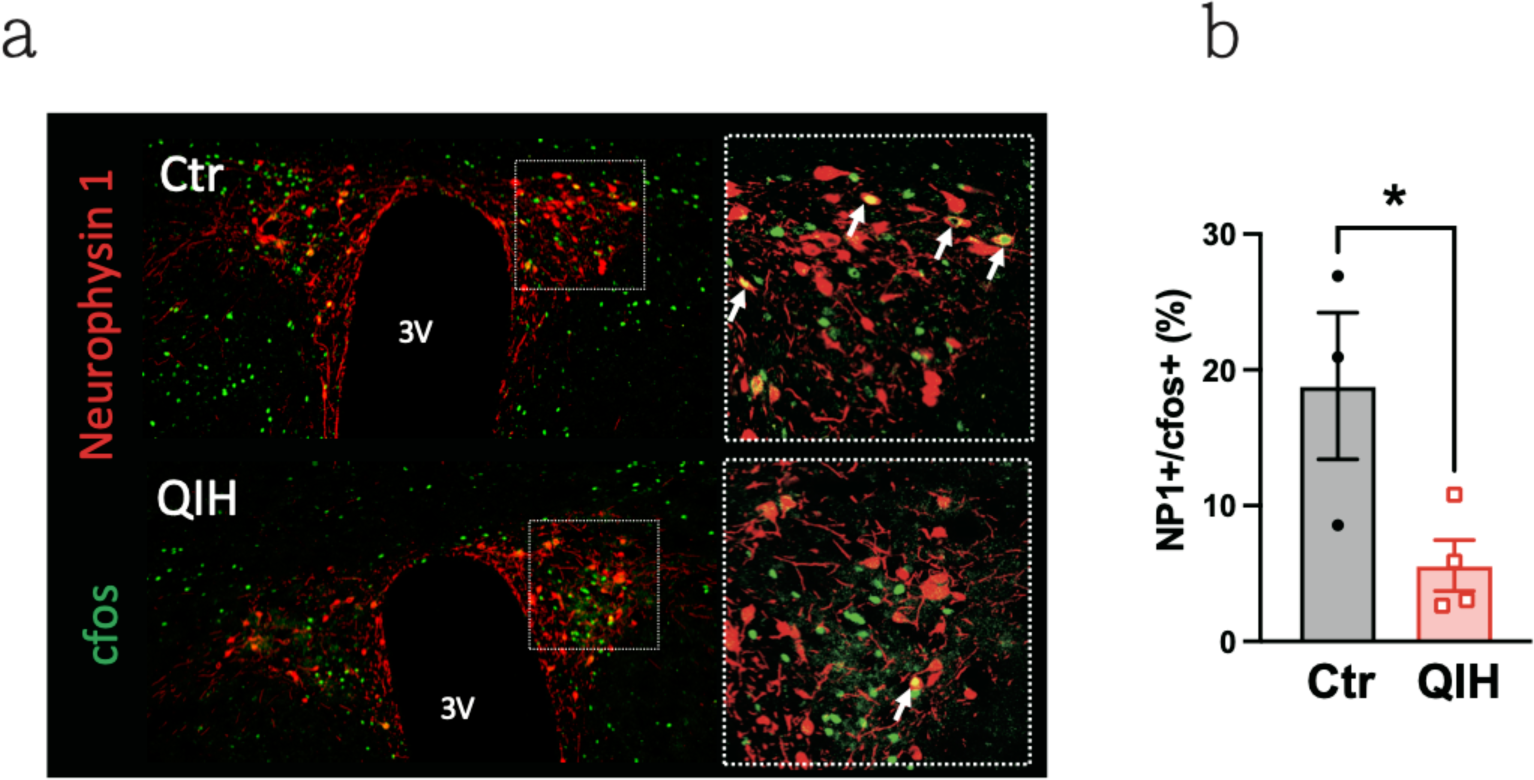

**Supplementary Figure 5.**
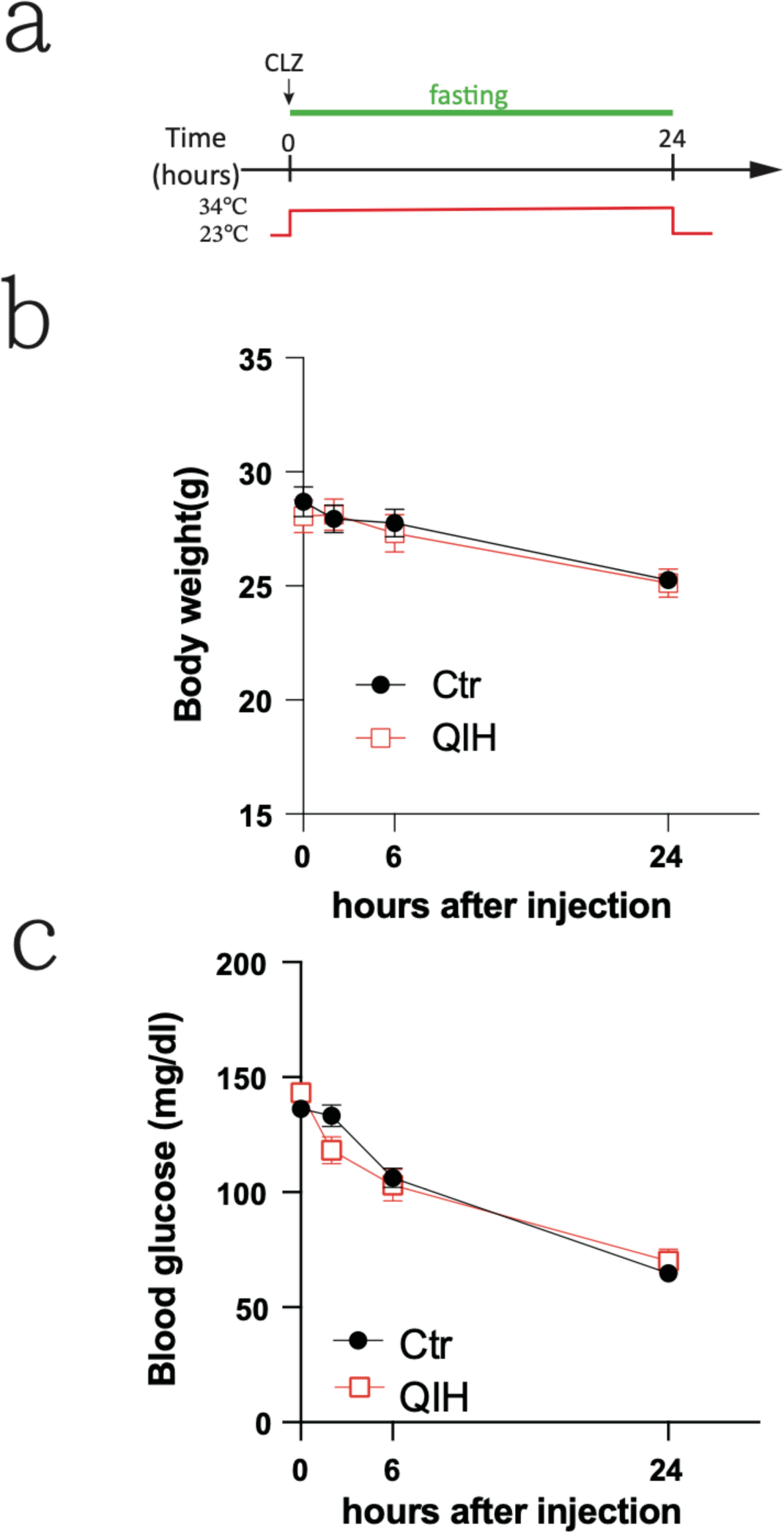

